# Distinct effects of tubulin isotype mutations on neurite growth in *Caenorhabditis elegans*

**DOI:** 10.1101/131326

**Authors:** Chaogu Zheng, Margarete Diaz-Cuadros, Ken C.Q. Nguyen, David H. Hall, Martin Chalfie

## Abstract

Tubulins, the building block of microtubules (MTs), play a critical role in both supporting and regulating neurite growth. Eukaryotic genomes contain multiple tubulin isotypes, and their missense mutations cause a range of neurodevelopmental defects. Using the *C. elegans* touch receptor neurons, we analyzed the effects of 67 tubulin missense mutations on neurite growth. Three types of mutations emerged: 1) loss-of-function mutations, which cause mild defects in neurite growth; 2) antimorphic mutations, which map to the GTP binding site and intradimer and interdimer interfaces, significantly reduce MT stability, and cause severe neurite growth defects; and 3) neomorphic mutations, which map to the exterior surface, increase MT stability, and cause ectopic neurite growth. Structure-function analysis reveals a causal relationship between tubulin structure and MT stability. This stability affects neuronal morphogenesis. As part of this analysis, we engineered several disease-associated human tubulin mutations into *C. elegans* genes and examined their impact on neuronal development at the cellular level. We also discovered an α-tubulin (TBA-7) that appears to destabilize MTs. Loss of TBA-7 led to the formation of hyperstable MTs and the generation of ectopic neurites; the lack of potential sites for polyamination and polyglutamination on TBA-7 may be responsible for this destabilization.

**Table of Content (TOC) Highlight Summary:** Different tubulin isotypes perform different functions in the regulation of MT structure and neurite growth, and missense mutations of tubulin genes have three types of distinct effects on MT stability and neurite growth. One α-tubulin isotype appears to induce relative instability due to the lack of potential post-translational modification sites.

## Introduction

Microtubules (MTs) play important roles in many aspects of neurite development, being involved in the formation, extension, guidance, and maintenance of neurites (reviewed in Dent *et al.*, 2011; Prokop, 2013; Sainath and Gallo, 2015). MTs constantly explore the growth cone periphery until they are captured by stabilized actin filaments or membrane receptors enriched at the side of the growth cone responding to a guidance cue (Tanaka *et al.*, 1995; Challacombe *et al.*, 1996; Schaefer *et al.*, 2008; Qu *et al.*, 2013). This capture transiently stabilizes MTs and enables MT elongation in the direction that the growth cone has turned. In addition to providing physical support for the neurite growth that follows the changes in actin dynamics, MTs also play an instructive role in neurite guidance. Since local application of the MT-stabilizing drug paclitaxel (also known as taxol) induced growth cone attraction, of the MT-destabilizing drug nocodazole induced repulsion, and of the MT-depolymerizing drug colchicine resulted in branching (Bray *et al.*, 1978; Buck and Zheng, 2002), signals that act by altering MT stability appear to directly initiate growth cone turning. Indeed, the guidance molecule Wnt can induce growth cone remodeling by changing the organization of MT structure through the inactivation of MT-plus end binding protein Adenomatous Polyposis Coli (Purro *et al.*, 2008). These results indicate that the regulation of MT dynamics is crucial for neurite growth and guidance, but how MT stability is controlled locally at specific sites of the growth cone and globally in the entire neuron is not well understood.

As the building blocks of MTs, α- and β-tubulins are crucial determinants of MT stability. In fact, eukaryotic genomes contain multiple tubulin genes encoding different isotypes (Sullivan, 1988; McKean *et al.*, 2001) that are expressed in spatially and temporally distinct patterns (Leandro-Garcia *et al.*, 2010) and that confer specific dynamic properties on MTs in tubulin polymerization assays *in vitro* (Hoyle and Raff, 1990; Panda *et al.*, 1994; Pucciarelli *et al.*, 2012). These observations support the “multi-tubulin hypothesis,” which proposes that distinct tubulin isotypes impart specific properties onto MTs, so they can perform particular cellular functions (Fulton and Simpson, 1976; Cleveland, 1987). Moreover, tubulins also undergo a range of complex post-translational modification (reviewed by Song and Brady, 2015), which affect the dynamics of MTs and their interaction with other proteins and can add to the complexity of the tubulin code. In neurons specifically, stable axonal MTs are more detyrosinated, acetylated, and glutamylated, whereas the dynamic MTs in the growth cone are more tyrosinated (Liao and Gundersen, 1998; Konishi and Setou, 2009). Tubulin isotypes and modifications at the C-terminal tail also control the velocity and processivity of MT motor proteins and microtubule depolymerization rates *in vitro* (Sirajuddin *et al.*, 2014). In turn MT motors can also regulate the organization of MTs (Verhey and Gaertig, 2007; Yu *et al.*, 2015).

The clinical importance of tubulin genes in the development of the nervous system is seen in the finding that α- and β-tubulin mutations in people are associated with microcephaly, lissencephaly, pachygyria, and other cortical malformations, as well as a range of axon guidance defects, including agenesis or hypoplasia of the corpus callosum, internal capsule, commissural fibers, and corticospinal tracts (Tischfield *et al.*, 2011). Three summaries of human mutations (Bahi-Buisson *et al.*, 2014; Liu and Dwyer, 2014; Chakraborti *et al.*, 2016) identify 60 point mutations in α-tubulin genes (51 in TUBA1A, 1 in TUBA3E, and 8 in TUBA4A), one splicing-affecting intronic deletion in TUBA8, 48 point mutations in β-tubulin genes (2 in TUBB2A, 24 in TUBB2B, 19 in TUBB3, and 3 in TUBB5), and one exonic deletion in TUBB2B that produce neurological disorders in heterozygous carriers (Table S1); all mutations, except for the TUBA8 deletion, appear to act as dominant-negative mutations. Although mutated residues are found throughout the molecules, many are found in region predicted to mediate either GTP binding, heterodimer stability, inter-dimer interaction, or association with motor proteins and other MAPs (Tischfield *et al.*, 2011). Despite some *in vitro* studies on the effects of the mutations on tubulin folding, heterodimer assembly, and MT growth (Jaglin *et al.*, 2009; Tian *et al.*, 2010; Tischfield *et al.*, 2010), very few *in vivo* studies can systematically test how these different tubulin mutations impact axon guidance and extension in living organisms. Moreover, the complexity of the mammalian nerve system makes the analysis of the developmental consequences of these tubulin mutations very difficult.

Here we use the morphologically simple and well-defined touch receptor neurons (TRNs) in the nematode *Caenorhabditis elegans* to model the effects of tubulin mutations on neurite growth. By analyzing a large collection of missense mutations in several tubulin genes, we found that these mutations caused three morphologically distinct defects in TRN neurite outgrowth: 1) the shortening of all TRN neurites; 2) the specific shortening of posteriorly directed neurites; and 3) the production of ectopic posteriorly directed neurites. The structural location of the mutated residue correlated with the resulting phenotype. Many tubulin mutations characterized in our study affect the same amino acid residue or region as the disease-causing mutations in humans. We generated several such human mutations in *C. elegans* tubulin genes through genome editing and found that they also caused distinct neurite growth defects that fall into the above categories. Thus, our system may be used to understand the different effects of the clinically identified tubulin mutations and to facilitate their classification. Moreover, we found that null mutations in two α-tubulin genes led to very different phenotypes, supporting the hypothesis that tubulin isotypes perform specific and often non-overlapping roles in neurite development.

## Results

### Neurite morphology and MT organization in the TRNs

The six mechanosensory TRNs (ALML/R, PLML/R, AVM, and PVM) in *C. elegans* are a useful model to study axonal outgrowth and guidance because of their well-defined morphology (Chalfie and Sulston, 1981). The ALM and PLM neurons are two pairs of embryonically derived, bilaterally symmetric cells, whereas the AVM and PVM neurons arise from postembryonic lineages. All six neurons have a long anteriorly-directed neurite (AN); in addition, the two PLM neurons have a posteriorly-directed neurite (PN), making them bipolar. Except in PVM, the ANs branch at their distal ends; we refer to this branch as the synaptic branch.

Unlike most of *C. elegans* the cells, which have only a few (∼5 per cross section in ventral cord neurons) 11-protofilament (11-p) MTs, TRNs contain a great number (∼31) of large diameter (15-protofilament, 15-p) MTs that assemble into bundles that fill the neurites (Chalfie and Thomson, 1979; Savage *et al.*, 1994). The abundance and prominence of the MTs in the TRNs permits easy observation of alterations in MT structure and organization by electron microscopy.

The *C. elegans* genome contains nine α-tubulin genes (*mec-12*, *tba-1*, *tba-2*, and *tba-4* through *tba-9*) and six β-tubulin genes (*ben-1*, *mec-7*, *tbb-1*, *tbb-2*, *tbb-4*, and *tbb-6*). Previous studies found that the α-tubulin MEC-12 and the β-tubulin MEC-7 are expressed specifically at high levels in the TRNs (Hamelin *et al.*, 1992; Mitani *et al.*, 1993) and are required for the mechanosensory functions of the TRNs (Chalfie and Au, 1989; Savage *et al.*, 1994). TRNs also express the ubiquitously present α-tubulin, TBA-1 and TBA-2, and β-tubulin, TBB-1 and TBB-2, but their loss had no effect on the development or function of TRNs (Fukushige *et al.*, 1993, 1995; Lockhead *et al.*, 2016). Here, we identify an additional α-tubulin TBA-6, which is also expressed in TRNs and functions to prevent excessive neurite growth.

In the following sections, we focus on MEC-7, MEC-12, and TBA-7. We first describe three types of *mec-7* missense mutations and their distinct effects on MT structure, neurite growth, and neuronal function, and then discuss the phenotypes of similar mutations in *mec-12*. These latter phenotypes are generally weaker than the corresponding *mec-7* mutations. To allow for a comparison of the *mec-7* and *mec-12* phenotypes, however, we have organized the first four figures by the type of data (Figure 1 for electron microscopy, Figure 2 for process outgrowth, Figure 3 for structural analysis, and Figure 4 for TRN activity). Finally, we describe the effect of the loss of *tba-7*.

**Figure 2.**
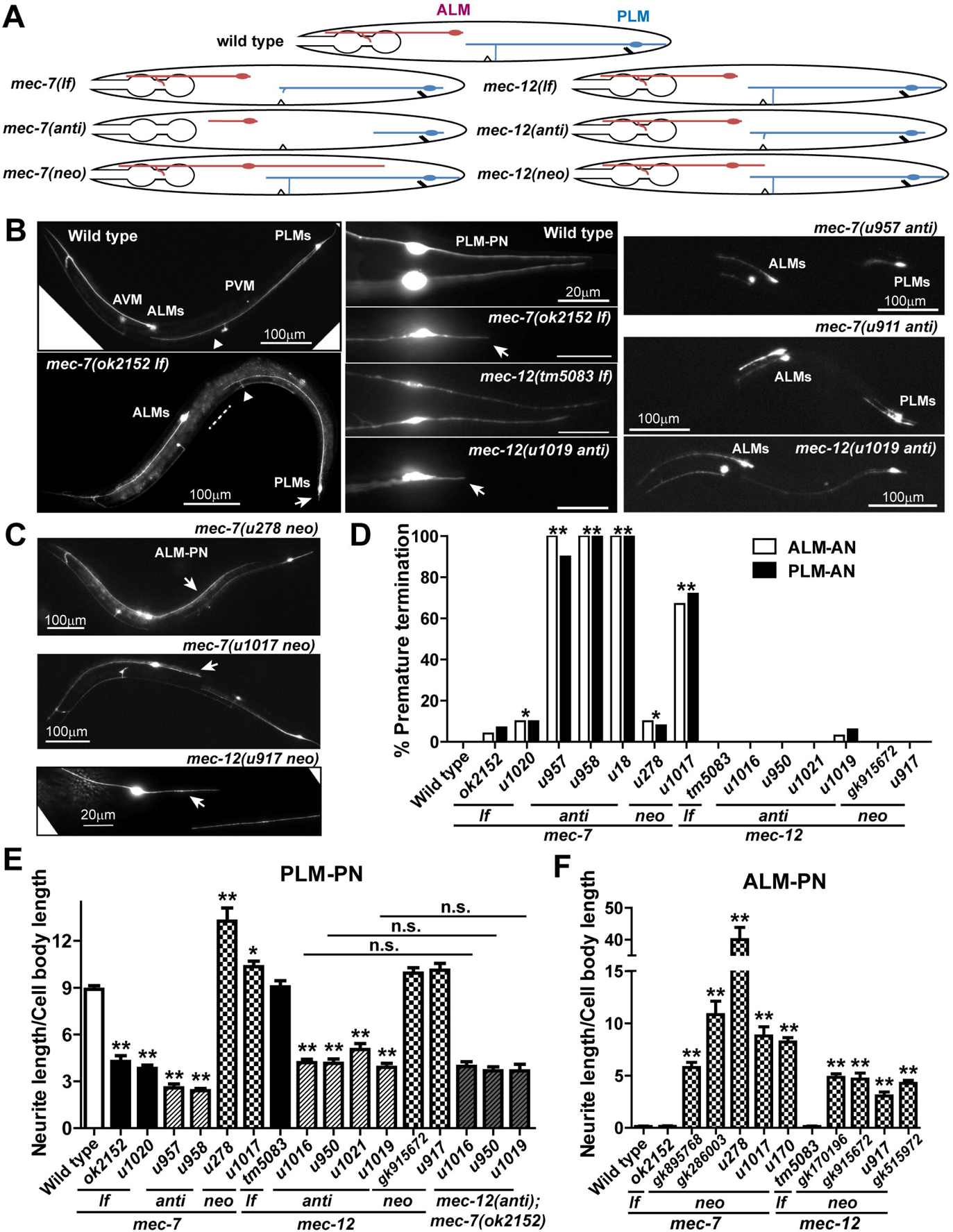
*mec-7* and *mec-12* mutations affect TRN neurite length. (A) Schematic diagram of ALM and PLM morphology in *mec-7* and *mec-12 lf*, *anti*, and *neo* mutants. (B) On the left panel, compared to the wild-type animals, *mec-7(ok2152 lf)* animals have an increased gap between PLM-AN and ALM cell body (dashed line) and significantly shortened PLM-PN (arrows). PLM-AN still extends beyond the position of the vulva indicated by the triangle. The middle panel shows PLM posterior neurites in the various animals. Arrows point to the shortened PLM-PN. The right panel displays the TRN morphologies in *mec-7[u957* (P171L) *anti]*, *mec-7[u958* (G244S) *anti]*, and *mec-12[u1019* (G354E) *anti]* animals. (C) TRN morphology of *mec-7[u278* (C303Y) *neo]*, *mec-7[u1017* (C377F) *neo]*, and *mec-12[u917* (V260I) *neo]* animals. Arrows point to the ectopic ALM-PN. (D) The percentage of ALM and PLM cells that had significantly shortened anterior neurites (ALM-AN not reaching the posterior pharyngeal bulb and PLM-AN not extending beyond the vulva). Amino acid changes in the alleles can be found in Table 1. A *χ*2 test for categorical data was used to compare mutants with the wild type. (E-F) The relative length (mean ± SEM) of PLM-PN or ALM-PN in *mec-7* and *mec-12 lf*, *anti*, and *neo* mutant animals, as well as the double mutants of *mec-7(ok2151 lf)* with *mec-12 neo* alleles. Dunnett’s tests were used to analyze the data.

**Figure 3.**
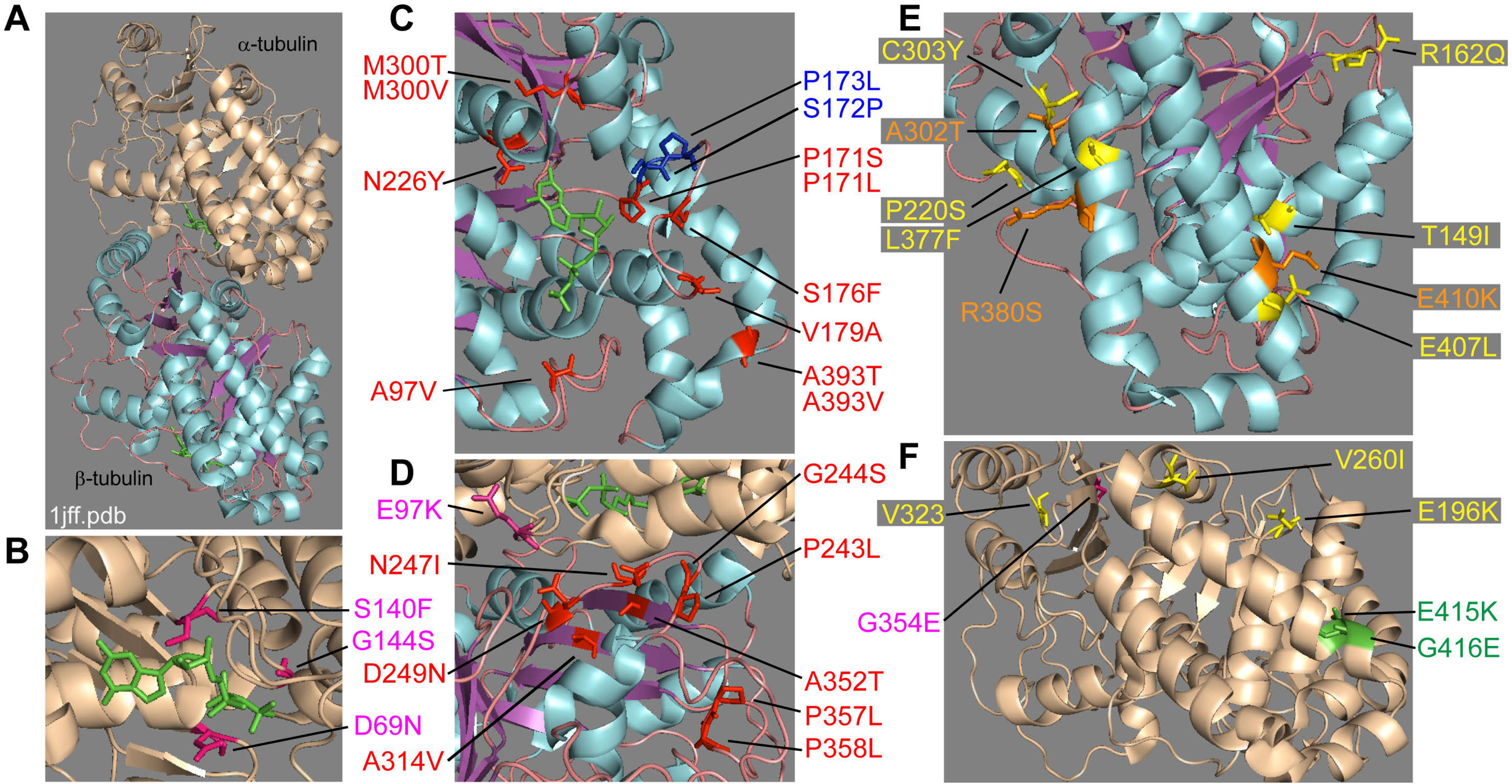
Position of amino acid residues changed in *mec-7* and *mec-12* mutants. (A) The structure of the *α /β* tubulin dimer (1jff.pdb) visualized using PyMOL. To be consistent with the original structure (Nogales *et al.*, 1998) and the mapping of human tubulin mutations (Tischfield *et al.*, 2011), α-tubulin (wheat) is shown on top of β-tubulin (cyan, magenta, and pink, which labels α-helices, β-sheets, and loops, respectively). GTP is labeled in green. (B-C) Amino acid changes around the GTP binding pocket in *mec-7* (red) and *mec-12* (pink) antimorphs. P173L and S172P (blue) are disease-associated mutations found in human TUBB3 that were engineered in *mec-7* to test their effects (see Figure 5). (D) *mec-7* (red) and *mec-12* (pink) *anti* mutations mapped to intradimer interface. (E-F) Amino acid alterations in *mec-7* (E, yellow) and *mec-12* (F, yellow) *neo* alleles. A302T, R380S, and E410K (orange) were clinically identified TUBB3 mutations. E415 and G416 (green), mutated in *mec-12* partial *lf* alleles, are located at the exterior surface of α-tubulin.

**Figure 4.**
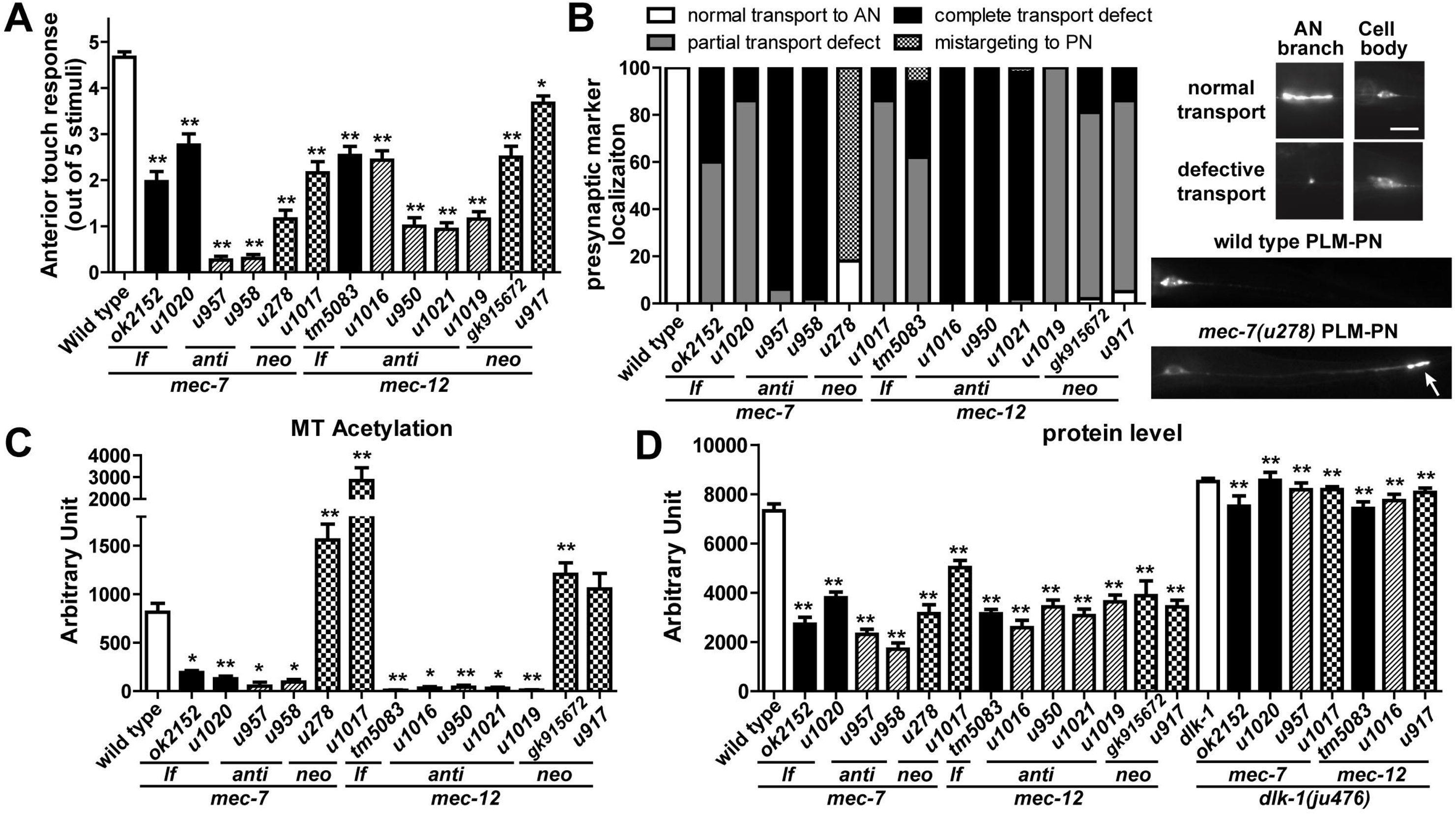
Effects of *mec-7* and *mec-12* mutations on TRN function and activity. (A) The average number of touch responses out of five for *mec-7* and *mec-12 lf*, *anti*, and *neo* mutants when they are touch anteriorly. (B) Defective targeting of the synaptic vesicle marker RAB-3::GFP in *mec-7* and *mec-12* mutant PLM neurons. RAB-3::GFP signal is found in patches where PLM-AN branches synapse onto target neurons in the wild-type ventral nerve cord. This targeting was partially (reduced GFP signal at the synapse) or completely (no GFP signal at the synapse) lost in many of the mutants; this loss in *mec-7(lf)* and *mec-12(anti)* mutants could be partly caused by the lack of PLM synaptic branch. In *mec-7(u278*; C303Y) mutants, however, the marker was mistargeted to the distal end of PLM-PN (arrow). (C) Immunofluorescent intensity of antibody labeling of acetylated α-tubulin. (D) Fluorescent intensity of the TRN marker *uIs134 [mec-17p::RFP]* in various *mec-7* and *mec-12* mutants and their doubles with the *dlk-1 lf (ju476)* allele. For (C-D), representative images are shown in Figure S3. Dunnett’s tests were performed to compare the mutants with the wild type animals, and *t* tests with Bonferroni correction were used to identify significant difference between the tubulin single mutants and *dlk-1; mec-7* or *dlk-1; mec-12* double mutants.

**Table 1.** The *mec-7* and *mec-12* mutations analyzed in this study. Several mutations are represented by multiple alleles, whose phenotypes were found to be similar. For touch sensitivity, + indicate the average response to 5 anterior stimuli is above 4; ± indicate the average is between 4 and 1; - indicate the average is below 1. Partial *lf* alleles of *mec-12* showed some but not all of the *lf* phenotypes (see the text). Asterisks indicate that the mutation was originally found in humans and was created in *mec-7* gene through CRISPR/Cas9-mediated genome editing. Mapping of the amino acid residues to the structural domains was done according to Tischfield *et al.* (2011).

| Gene | Allele | Mutation | Structural function | Classification | Morphological defects | Touch sensitivity | Expression |
| --- | --- | --- | --- | --- | --- | --- | --- |
| *mec-7* | *u319* | S25F | Tubulin folding | weak antimorph | moderately shortened neurites | - | semidominant |
|  | *u305, u1020* | G34S | Lumen-facing loop | lf | short PLM-PN | ± | recessive |
|  | *gk906464* | G38E | Lumen-facing loop | N/A | no defects | + | N/A |
|  | *u58, u223* | P61L | Lateral interaction | weak antimorph | moderately shortened neurites | - | semidominant |
|  | *u249* | P61S | Lateral interaction | weak antimorph | moderately shortened neurites | - | semidominant |
|  | *gk337318* | V64I | Tubulin folding | N/A | no defects | + | N/A |
|  | *u430* | A97V | GTP binding | antimorph | short TRN neurites | - | recessive |
|  | *u222* | G109E | Lateral interaction | weak antimorph | moderately shortened neurites | - | semidominant |
|  | *u429, u433* | G141E | Tubulin folding | lf | short PLM-PN | - | recessive |
|  | *gk595364* | G141R | Tubulin folding | lf | short PLM-PN | - | recessive |
|  | *u275* | G148R | Tubulin folding | lf | short PLM-PN | - | recessive |
|  | *gk895768* | T149I | Tubulin folding | neomorph | ectopic ALM-PN | ± | recessive |
|  | *gk286003* | R162Q | MAP binding | neomorph | ectopic ALM-PN | ± | recessive |
|  | *u911* | P171S | GTP binding | antimorph | short TRN neurites | - | dominant |
|  | *u957, u127, e1343* | P171L | GTP binding | antimorph | short TRN neurites | - | semidominant |
|  | *u1056* | S172P* | GTP binding | lf | short PLM-PN | - | recessive |
|  | *u1057* | P173L* | GTP binding | N/A | no defects | + | N/A |
|  | *u48* | S176F | GTP binding | antimorph | short TRN neurites | - | semidominant |
|  | *u449* | V179A | GTP binding | antimorph | short TRN neurites | - | semidominant |
|  | *u10* | S188F | Tubulin folding | lf | short PLM-PN | ± | recessive |
|  | *u225* | T214P | Tubulin folding | lf | short PLM-PN | ± | recessive |
|  | *ky852* | P220S | Tubulin folding | neomorph | ectopic ALM-PN | - | semidominant |
|  | *u262* | N226Y | GTP binding | antimorph | short TRN neurites | - | semidominant |
|  | *gk286002* | P243S | Intradimer interaction | lf | short PLM-PN | ± | recessive |
|  | *u283* | P243L | Intradimer interaction | antimorph | short TRN neurites | - | dominant |
|  | *u129, u958* | G244S | Intradimer interaction | antimorph | short TRN neurites | - | dominant |
|  | *n434* | N247I | Intradimer interaction | antimorph | short TRN neurites | - | dominant |
|  | *u162* | D249N | Intradimer interaction | antimorph | short TRN neurites | - | dominant |
|  | *e1505* | G269D | Tubulin folding | lf | short PLM-PN | ± | recessive |
|  | *e1527* | V286D | Lateral interaction | antimorph | short TRN neurites | - | dominant |
|  | *u445* | M300V | Tubulin folding | antimorph | short TRN neurites | - | semidominant |
|  | *u98* | M300T | Tubulin folding | antimorph | short TRN neurites | - | semidominant |
|  | *u1058* | A302T* | MAP binding | neomorph | ectopic ALM-PN | ± | semidominant |
|  | *u278* | C303Y | MAP binding | neomorph | ectopic ALM-PN | - | semidominant |
|  | *gk286001* | C303S | MAP binding | N/A | no defects | + | N/A |
|  | *gk286000* | A314V | Tubulin folding | antimorph | short TRN neurites | - | semidominant |
|  | *e1522* | F317I | Tubulin folding | lf | short PLM-PN | ± | recessive |
|  | *u234* | R318Q | Tubulin folding | lf | short PLM-PN | ± | recessive |
|  | *u955* | A352T | Intradimer interaction | antimorph | short TRN neurites | - | dominant |
|  | *u910, gk373602* | P357L | Tubulin folding | antimorph | short TRN neurites | - | dominant |
|  | *u956* | P358L | Tubulin folding | antimorph | short TRN neurites | – | dominant |
|  | *u428* | G369E | Tubulin folding | lf | short PLM-PN | – | recessive |
|  | *u1017* | L377F | MAP binding | neomorph | the growth of ectopic ALM-PN | ± | recessive |
|  | *u1059* | R380S* | MAP binding | neomorph | the growth of ectopic ALM-PN | ± | recessive |
|  | *u18* | A393T | Longitudinal interaction | antimorph | short TRN neurites | – | dominant |
|  | *gk285997* | A393V | Longitudinal interaction | antimorph | short TRN neurites | – | dominant |
|  | *u170* | E407L | MAP binding | neomorph | the growth of ectopic ALM-PN | ± | recessive |
|  | *u1060* | E410K* | MAP binding | antimorph | shortened TRN-ANs | – | semidominant |
| *mec-12* | *gk170196* | P32S | Lumen-facing loop | neomorph | the growth of ectopic ALM-PN | ± | recessive |
|  | *gk170195* | S50N | Lumen-facing loop | N/A | no defects | + | N/A |
|  | *gk636747* | R60H | Lumen-facing loop | N/A | no defects | + | N/A |
|  | *u76* | D69N | GTP binding | antimorph | short PLM-PN | – | recessive |
|  | *u1016* | E97K | Intradimer interaction | antimorph | short PLM-PN | – | recessive |
|  | *u950, gk672907* | S140F | GTP binding | antimorph | short PLM-PN | – | recessive |
|  | *gk600523* | G142E | GTP binding | lf | no defects | ± | recessive |
|  | *u1021, e1607* | G144S | GTP binding | antimorph | short PLM-PN | – | recessive |
|  | *gk583647* | L152F | Tubulin folding | N/A | no defects | + | N/A |
|  | *u50, e1605* | H192Y | MAP binding | partial lf | no defects | – | recessive |
|  | *gk915672* | E196K | MAP binding | neomorph | the growth of ectopic ALM-PN | ± | recessive |
|  | *u1041* | G246E | Tubulin folding | lf | no defects | ± | recessive |
|  | *u917* | V260I | MAP binding | neomorph | the growth of ectopic ALM-PN | ± | recessive |
|  | *gk854211* | P307L | MAP binding | N/A | no defects | + | N/A |
|  | *gk515972* | V323I | Longitudinal interaction | neomorph | the growth of ectopic ALM-PN | ± | recessive |
|  | *u241, u1019* | G354E | Longitudinal interaction | antimorph | short PLM-PN | – | recessive |
|  | *gk341552* | G365E | Lumen-facing loop | N/A | no defects | + | N/A |
|  | *u63* | E415K | MAP binding | partial lf | no defects | ± | recessive |
|  | *gm379* | G416E | MAP binding | partial lf | no defects | ± | recessive |

**Figure 1.**
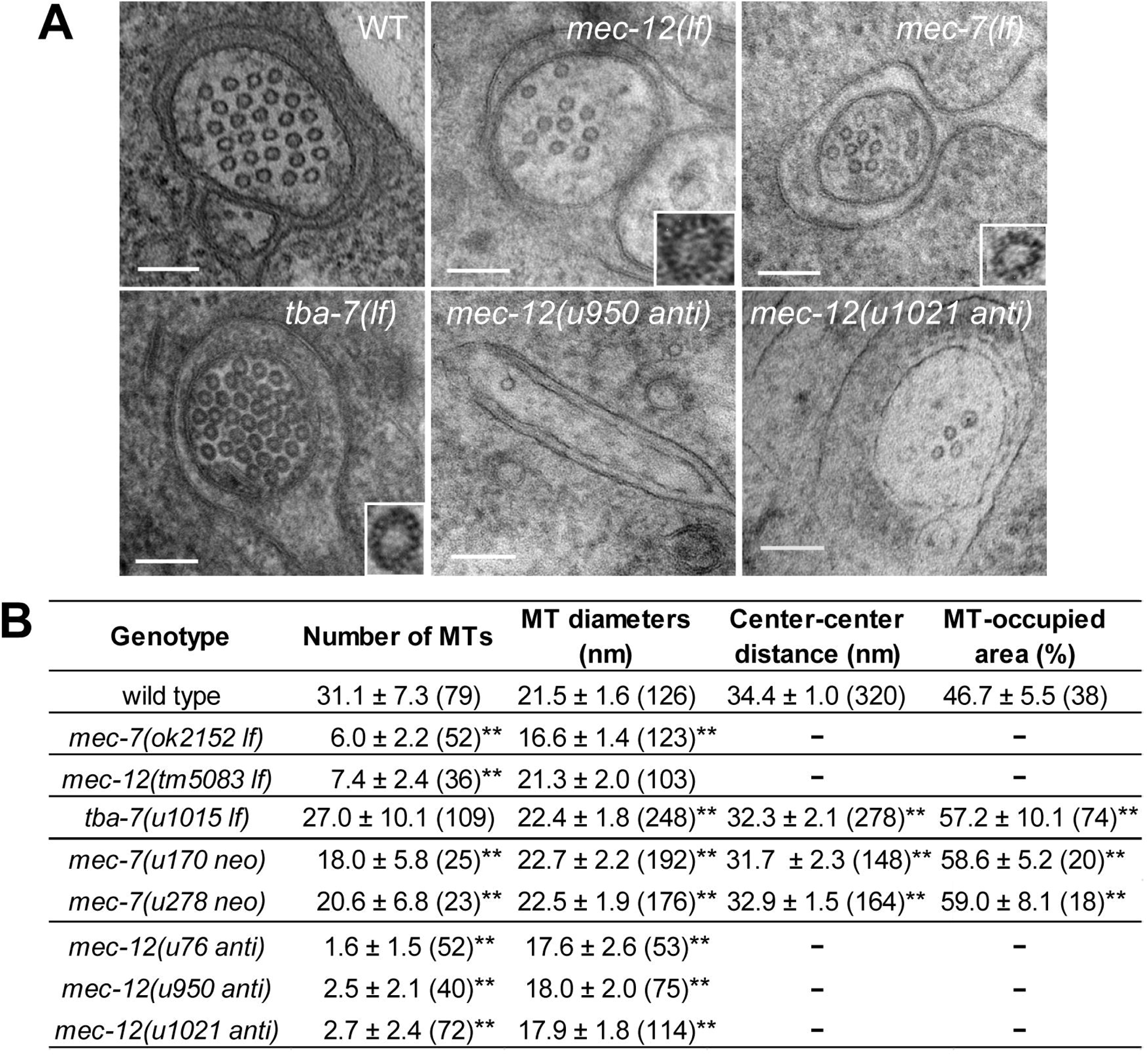
MT structures in tubulin mutants. (A) Cross sectional images of ALM-AN in wild type and *mec-12(tm5083 lf)*, *mec-7(ok2152 lf)*, *tba-7(u1015 lf) mec-12[u950* (S140F) *anti]* and *mec-12[u1021* (G144S) *anti]* mutants. Images for *mec-7(anti)* and *mec-7(neo)* can be found in previous publications (Chalfie and Thomson, 1982; Savage *et al.*, 1994). Insets in the lower right corner show the 15-p, 11-p, and 15-p structure of tannic acid stained MTs of *mec-12(lf)*, *mec-7(lf)*, and *tba-7(lf)* animals, respectively (4-fold enlarged). Scale bar = 100 nm. (B) Measures of MT structure and organization. Mean ± SD are shown, and numbers of observations are in parentheses. A Dunnett’s test was performed to compare the mutants with the wild type. Throughout the figures, one asterisk represents a statistical significance of *p* < 0.05 and two asterisks indicate *p* < 0.01. We did not measure center-center distance and MT-occupied area in *mec-7(lf)*, *mec-12(lf)*, and *mec-12(anti)* mutants, because they contained very few MTs, which did not form bundles.

### Deletion of *mec-7*/β-tubulin led to the loss of 15-p MTs and defects in posteriorly directed neurite growth

Using electron microscopy (EM), we found that *mec-7(ok2152)* knockout animals had on average 6 MTs in a cross section of the ALM neurite compared to 31 MTs in the wild type animals (Figure 1). The *ok2152* mutation deletes the first four exons (Figure S1A) and is thus is a likely null allele. The diameter of MTs in *mec-7(ok2152)* animals was much smaller (16.6 nm) than that of the wild type (21.5 nm), and tannic acid staining indicated that the 15-p MTs in TRNs were replaced by smaller 11-p MTs in the absence of MEC-7 (Figure 1). These results suggest that the abundance of MTs and the formation of large diameter 15-p MTs requires MEC-7, which is consistent with our previous observations on *mec-7(e1506)* animals (the *e1506* mutation alters the start codon and results in no detectable *mec-7* mRNA; Chalfie and Thomson, 1982; Savage *et al.*, 1994).

We next examined how *mec-7*(*ok2152*) mutation affected TRN morphogenesis. The *mec-7* deletion allele *mec-7* only slightly affected the growth of anteriorly-directed neurites of PLM neurons but markedly shortened the posteriorly-directed neurites (Figure 2B). This result suggests that PLM-AN and PLM-PN, which arise at the same time in the embryo, have different tubulin requirements. Normally the PLM-AN extends anteriorly past the vulva to within 50 μm of ALM cell body. Lockhead *et al.* (2016) reported that the gap between the end of PLM and ALM cell body was wider in animals with the presumed null *mec-7(u241)* due to the shortening of the PLM-AN. We found a similar modest widening of the gap in *mec-7(ok2152)* animals, but 92% of the PLM-ANs still extended beyond the vulva (Figure 2B and D). Moreover, no shortening of the ALM-AN was seen. A more penetrant and striking defect, however, was the loss of the synaptic branch in 78% of PLM-ANs in *mec-7(ok2152)* animals (Figure 2A and S2; the synaptic branch normally arises posterior to the vulva). The ALM synaptic branch was not affected. *mec-7* nonsense alleles *u156* and *u440* produced similar morphological defects (Figure S1B and data not shown). These results suggest that the smaller, generic 11-p MTs in *mec-7* knockout mutants are, in general, capable of supporting the growth of most TRN neurites, but the normal extension of PLM-PN and the elaboration of the posterior synaptic branch require the special 15-p MTs.

The neurites of wild-type TRNs have many 15-p MTs associated into a bundle. This bundle may provide stronger support for neurite growth than the few 11-p MTs found in *mec-7* deletion mutants. If so, the requirement for the 15-p MTs in the PLM-PN may be due to a higher sensitivity to changes in microtubule stability there than in the PLM-AN. In fact, treatment with the MT-destabilizing drug colchicine (1mM), which specifically removes most of the TRN MTs (Chalfie and Thomson, 1982), led to a similar shortening of the PLM-PN, phenocopying the *mec-7* null mutants (Figure S3A).

### Antimorphic mutations in *mec-7* led to severe neurite growth defects

We next examined 55 *mec-7* alleles that represent 48 different missense mutations (Table 1) and found three categories of TRN morphological defects. First, 12 mutations caused neurite growth defects similar to the knockout allele and were thus classified as loss-of-function (*lf*) mutations. Data from *u1020* (G34S) animals is shown in Figure 2 as an additional *lf* example. When mapped to the structure of a bovine αβ tubulin dimer (Figure 3; Nogales *et al.*, 1998), *lf* mutations were found throughout the molecule and presumably affected protein folding. Second, 19 mutations caused severe shortening of all TRN neurites; the ALM-AN did not reach the pharynx, the PLM-AN terminated before reaching the PVM cell body and the PLM-PN was significantly shortened (Figure 2B and D). Because of the dominant-negative nature of these alleles (all, except *u430*, were either dominant or semi-dominant), we classified them as antimorphic (*anti*) gain-of-function mutations. We also identified 4 weak *mec-7(anti)* mutants (Table 1), in which ALM-AN did not extend beyond the nerve ring and PLM-AN did not reach the vulva. Third, 6 mutations led to the growth of an extra posteriorly-directed neurite in ALM neurons, as well as the overextension of the PLM-PN. Among the six alleles, *u278* (C303Y) produced the strongest phenotype; the ectopic posterior neurite of ALM neurons often extended posteriorly to the PLM cell body (Figure 2C). Because this phenotype was novel and different from the phenotype of either *mec-7 lf* or *anti* mutations, we classified these alleles as neomorphic (*neo*) gain-of-function mutations. Except two semi-dominant ones (*u278* and *ky852*), most of the *neo* alleles were recessive. Because of the antagonism between anteriorly and posteriorly directed outgrowth (Zheng *et al.*, 2016), some *neo* alleles, especially *u1017*, caused mild shortening of the ANs, i.e. PLM-AN terminated slightly posterior to the vulva (Figure 3C).

EM studies of the *mec-7(anti)* allele, *e1343* (P171L), found that the mutants contained very few (2.8 ± 0.5 in a cross section) MTs (Chalfie and Thomson, 1982), which suggests that the *anti* mutations block MT polymerization and thereby cause the severe neurite outgrowth defects. In fact, when mapped to the tubulin structure, amino acid residues mutated in the *anti* alleles are clustered in three regions: 1) the GTP/GDP binding pocket, 2) the intradimer interface, and 3) the lateral or longitudinal interdimer interface. All of which are important for tubulin polymerization. For example, *u430* (A97V), *u911* (P171S), *u957* (P171L), and *u262* (N226Y) mutations alter amino acids in direct contact with the GTP molecule, and *u48* (S176F) and *u449* (V179A) change amino acids in the B5 (the fifth β-strand)-to-H5 (the fifth α-helix) loop, which is crucial for forming the GTP/GDP binding pocket (Figure 3C). Two other mutations *u445* (M300V) and *u98* (M300T) substitute M300, which is located in a loop near the GTP binding site and may participate in positioning H7 (the seventh α-helix) with its critical N226, and thus indirectly affect GTP binding.

The second group of mutated residues in *mec-7(anti)* alleles is located in the intradimer interface, where β-tubulin makes contact with α-tubulin to form the heterodimer (red residues in Figure 3D). *u283* (P243L), *u958* (G244S), *n434* (N247I), and *u162* (D249N) mutations all affect the residues on the H7-to-H8 loop, which interacts extensively with residues on H1 and H2 of α-tubulin. Two other dominant mutations *gk286000* (A314V) and *u955* (A352T) changed residues that located on B8 and B9, respectively, which are physically adjacent to the H7-to-H8 loop at the interface. In addition, two prolines mutations, *u910* (P357L) and *u956* (P358L) possibly disrupted the B9-to-B10 loop (a.a. 356-361) critical for the positioning of B9.

The third group of *mec-7(anti)* alleles include *e1527* (V286D), *u18* (A393T), and *gk285997* (A393V), which affect residues involved in the lateral (V286) and longitudinal (A393) interdimer interaction between one α/β-tubulin heterodimer and the neighboring one on the MTs. These mutations may interfere with the interaction between tubulin dimers.

Overall, the MEC-7/β-tubulin *anti* mutants likely act in a dominant-negative manner by forming poisonous α/β dimers, whose incorporation into MTs could terminate MT polymerization and induce instability. The mutated β-tubulin either cannot properly bind to GTP (the first group), form misshaped α/β heterodimers that block the growing end of MTs (the second group) or disrupt the stacking of tubulin dimers (the third group). Therefore, changes in tubulin structure led to compromised MT elongation, which caused severe defects in neurite outgrowth, highlighting the importance of functional MTs in neuronal morphogenesis.

### Neomorphic mutations in *mec-7* led to ectopic neurite growth by increasing MT stability

The *mec-7(neo)* mutations appear to induce the growth of ectopic ALM-PNs by forming hyperstable MTs, as first suggested by Kirszenblat *et al.* (2013). Supporting this hypothesis, pharmacological stabilization of MTs with paclitaxel induced the growth of ALM-PN, although at a low frequency, and destabilizing MTs with colchicine can partially suppress this ectopic growth in *mec-7(ky852 neo)* mutants (Kirszenblat *et al.*, 2013) and in *mec-7(u278 neo)* and *mec-7(u1017 neo)* animals (Figure S3A). We found that *mec-7(neo)* animals had increased resistance to colchicine compared to the wild-type animals. More of this MT-destabilizing drug was needed to produce the same level of reduction in both TRN function (touch sensitivity) and PLM-PN length in *neo* mutants than in wild-type animals (Figure S3B-C). Moreover, these mutants also showed increased tubulin acetylation (Figure 4C), an indication of stable MTs (Song and Brady, 2015).

The greater MT stability and higher resistance to destabilizing reagents in the *neo* mutants could be the result of either reduced dynamics of individual MTs or more stable bundles. MTs in wild-type TRNs showed very little dynamics, since Hsu *et al.* (2014) found and we confirmed that the GFP labeled MT plus-end binding protein EBP-2/EB2 barely moved in the wild-type TRNs. Thus, the already very high stability of individual MTs is unlikely to increase greatly in the *neo* mutants. However, the MT bundles in the TRNs of *neo* mutants have changed and so may be the cause of the drug-resistance in these animals. By quantifying EM data previously obtained (Savage *et al.*, 1994), we found that *mec-7(u278 neo)* and *mec-7(u170 neo)* mutants kept the large diameter MTs (slightly larger than in wild-type cells), but had fewer of them (Figure 1B). Importantly, the MT bundles in the mutants are more closely spaced than the wild type and are not surrounded by the electron-lucent region seen in the wild type (Savage *et al.*, 1994). We quantified these differences by showing that the distance between two closest center points of MTs is smaller in *mec-7(neo)* mutants compared to the wild type and that MTs occupied a bigger proportion of the cross-sectional area of the neurite in the mutants (Figure 1B). In fact, the TRN neurite in these mutants is about 50% thinner than the wild type animals. The observation that tightly packed 15-p MT bundles filled up most of the space in the TRN neurites is consistent with increased MT stability, which presumably allowed the excessive neurite growth towards the posterior, overcoming the normal inhibition of this growth in the wild-type ALM neurons.

The majority of the altered residues in the *mec-7(neo)* alleles are located on the exterior surface of MTs, where they may interfere with the binding of motor proteins or MAPs to the MTs and therefore cause changes in MT stability. Examples include *u1017* (L377F) and *u170* (E407K), which are located on the H11 and H12 helices, respectively, *gk286003* (R162Q) on the H4-to-B5 loop, and *u278* (C303Y) on the H9-to-B8 loop are also exposed on the MT surface (Figure 3E). Interestingly, although *u278* (C303Y) produced the strongest *neo* phenotype, C303S mutation in allele *gk286001* did not lead to similar excessive growth (Table 1), suggesting that tyrosine’s bulky phenolic ring may be responsible for the phenotype. In contrast, the *neo* alleles *gk895768* (T149I) and *ky852* (P220S) affect residues on the inside the MTs and may be important for the folding of the protein.

### Loss-of-function, antimophic, and neomorphic mutations of *mec-12*/α-tubulin caused similar but weaker phenotypes than the *mec-7* mutations

We next examined the contribution of MEC-12/α-tubulin to MT organization and neurite growth in the TRNs. *mec-12(tm5083)* allele deletes part of exon 3 and intron 3 (Figure S1B) and is likely to be a *lf* mutation. Those deletion mutants had much fewer (∼7) MTs in a cross section of ALM neurites than the wild type animals, but the MTs had the same diameter as the 15-p MTs in the wild type (Figure 1B). Moreover, we observed no defects in TRN outgrowth in the *mec-12(tm5083 lf)* mutants (Figure 2) and confirmed the results using three CRISPR/Cas9-induced frameshift mutations (*u1026*, *u1027*, and *u1028*; Figure S1B and data not shown). Therefore, although the abundance of MTs requires MEC-12, the formation of 15-p MTs does not require MEC-12, and the reduction of MT numbers in *mec-12(lf)* mutants did not affect neurite development. Because *C. elegans* genome contains nine α-tubulin genes, other tubulin isotypes may compensate for the loss of MEC-12. Although TBA-1 and TBA-2 are expressed in the TRNs, *tba-1(lf)*; *mec-12(lf)* and *tba-2(lf); mec-12(lf)* double mutants had very mild defects in neurite growth, only the slight shortening of PLM-AN (Lockhead *et al.*, 2016). In this study, we found another α-tubulin TBA-7, which is also expressed and functions in TRNs (see below); however, *mec-12; tba-7* double mutants were still capable of forming and growing normal TRN neurites, suggesting further genetic redundancy.

The EM results of *mec-12(lf)* mutants were unexpected, because the *mec-12[(e1607* (G144S)] allele, which had been regarded as the null allele (Bounoutas *et al.*, 2009a; Bounoutas *et al.*, 2011; Hsu *et al.*, 2014), had very few (∼2) MTs, all of which had the small diameter (Chalfie and Au, 1989). We repeated the EM studies using the newly isolated *mec-12(u1021)* allele that carries the same G144S mutation and found the ALM neurites had ∼2.7 MTs in a cross section and 15% of the 72 sections examined had no MTs; the diameter of MTs was 17.9 nm (Figure 1B). We found that *e1607* and several other missense alleles (Table 1) were in fact antimorphic (*anti*) gain-of-function alleles, which caused the significant shortening of PLM-PN (Figure 2B and E). Like the *mec-7(anti)*, the five *mec-12(anti)* mutations also mapped to the GTP binding pocket [*u76* (D69N), *u950* (S140F), and *u1021* (G144S); labeled in red in Figure 3B], the intradimer interface [*u1016* (E97K); Figure 3D], or the interdimer interface [*u1019* (G354E)]. However, unlike the *mec-7(anti)* alleles, *mec-12(anti)* are mostly recessive at 20 °C, except for *u1019*, and did not cause strong, general defects in TRN neurite outgrowth, but rather specific shortening of PLM-PN similar to the *mec-7(lf)* mutants.

One possible explanation for this phenotype is that MEC-12/α-tubulin preferentially or exclusively binds to MEC-7/β-tubulin, and the MEC-12(*anti*) proteins sequester and disable MEC-7 but no other β-tubulin isotypes. As a result, both *mec-12(anti)* and *mec-7(lf)* removes MEC-7 from the pool of free tubulins, leading to the loss of the 15-p MTs that require MEC-7 and the shortening of PLM-PN. Consistent with this hypothesis, *mec-7(lf); mec-12(anti)* double mutants did not show additive effects with regards to the neurite growth defects compared to either single mutant (Figure 2E).

Four of the 19 *mec-12* missense mutations (Table 1) were *neo* alleles [*gk170196* (P32S), *gk915672* (E196K), *u917* (V260I), *gk515972* (V323I)]. These alleles produced a similar but less severe phenotypes than the *mec-7(neo)* alleles. Specifically, the ectopic ALM-PN was shorter in the *mec-12(neo)* mutants than in the *mec-7(neo)* mutants (Figure 2F). This observation supports the notion that MEC-12 may be less important than MEC-7 in regulating MT stability and subsequent neurite growth. E196 and V260 are located on loops on the exterior surface, whereas P32 and V323 may be involved in protein folding (Figure 3F).

The production of the ectopic ALM-PN requires both *mec-7* and *mec-12*. The deletion of *mec-12* suppressed the growth of ALM-PN in *mec-7(neo)* mutants, and the deletion of *mec-7* similarly suppressed the effect of *mec-12(neo)* alleles (Figure S3A). These results suggest that the tubulin *neo* mutations induce excessive neurite growth by altering the properties of mostly MEC-7/MEC-12 heterodimers and not other α/β heterodimers.

### *mec-7* and *mec-12* mutations also affect TRN functions, vesicle transport, and protein expression

In addition to neurite growth, MTs are important for many other functions in TRNs (Bounoutas *et al.*, 2009a; Hsu *et al.*, 2014). Deletion of either *mec-12* or *mec-7* resulted in touch insensitivity, defects in the localization of pre-synaptic vesicles, and severe loss of MT acetylation, a mark for stable MTs (Figure 4 and S4A). Since MEC-12 is the only α-tubulin isotype that contains the acetylation site (lysine 40) in *C. elegans*, the TRN-specific 15-p MTs are likely the only MTs that can be acetylated. The loss of *mec-7* and *mec-12* also induced a global reduction in protein levels through a mechanism dependent on the MAPK pathway dual leucine zipper-bearing kinase DLK-1 (Bounoutas *et al.*, 2011). Interestingly, mutations in *dlk-1* restored the normal protein expression in *mec-7* and *mec-12* null mutants but failed to rescue the PLM-PN growth defects in *mec-7(lf)* animals (Figure 4D and S4B). Therefore, this PLM-PN defect is most likely a direct effect of the loss of MEC-7 and not the result of secondary changes in protein levels.

Compared to the *mec-7(lf)* mutants, the *mec-7(anti)* mutations caused stronger defects not only in neurite outgrowth, but also in mechanosensation, synaptic vesicle transport, tubulin acetylation, and protein expression levels (Figure 4 and S4B). Those phenotypes in *mec-12(anti)* mutants were more severe than the phenotypes in the *mec-12(lf)* alleles and were comparable to those produced by *mec-7(lf)* mutations but not as strong as the *mec-7(anti)* mutations (Figure 4). The *mec-7* and *mec-12 neo* alleles also caused touch insensitivity, synaptic vesicle transport defects, and reduced TRN protein levels, as found in the *lf* mutants; but the *neo* mutants had markedly increased MT acetylation (Figure 4). Moreover, the fact that both impaired (in *anti* mutants) and excessive (in *neo* mutants) neurite growth can occur in conjunction with other TRN defects suggests that the role of MTs in regulating neurite development is genetically separable from other MT functions.

Furthermore, several *mec-12* partial *lf* mutations affected only a subset of MT functions. *mec-12(e1605)* animals carrying the H192Y missense mutation were defective in mechanosensation but had retained normal 15-p MT structures, tubulin acetylation, axonal transport, protein levels, and neurite growth patterns (Figure S5; Bounoutas *et al.*, 2009a).

Similarly, *mec-12* mutants carrying the *u63* (E415K) or *gm379* (G416E) alleles were partially touch-insensitive but kept the large dimeter MTs; however, these animals showed mistargeting of synaptic vesicles presumably because alterations in the EEGE (amino acid 414-417) motif increased affinity with the motor dynein (Hsu *et al.*, 2014). *u63* and *gm379* mutations also caused a slight decrease in tubulin acetylation and a partial reduction in protein production; *gm379* allele led to very mild defects in the growth of PLM-PN (Figure S5). These mutations may be useful in understanding some specific aspects of MT functions.

### Modeling the effects of tubulin mutations using the TRN neurites

Our genetic analysis shows that missense mutations in tubulin genes can have distinct effects on MT stability and neurite growth pattern, and those different phenotypes appear to correlate with the positions of the altered residues in the tubulin structure. Such structure-function analyses have been carried out for the yeast α-tubulin TUB1 and the Drosophila testis-specific β-tubulin βTub85D (Fackenthal *et al.*, 1995; Richards *et al.*, 2000) but not for tubulins expressed in the nervous system. Moreover, recent discovery of over a hundred missense mutations in tubulin genes in patients with a range of neurological disorders prompted us to examine the cellular impact of specific tubulin mutations on neurons. Combining the convenience of genome editing in *C. elegans* and the ease of observing TRN morphology as the readout of MT stability and neurite growth, we could systematically study the effects of those human tubulin mutations. To provide a proof of concept, we generated five clinically observed β-tubulin mutations (S172P, P173L, A302T, P380S, and E410K) in *mec-7* through CRISPR/Cas9-mediated gene editing (Materials and Methods; Figure 5A) and found that these mutations indeed caused distinct phenotypes on TRN morphogenesis.

**Figure 5.**
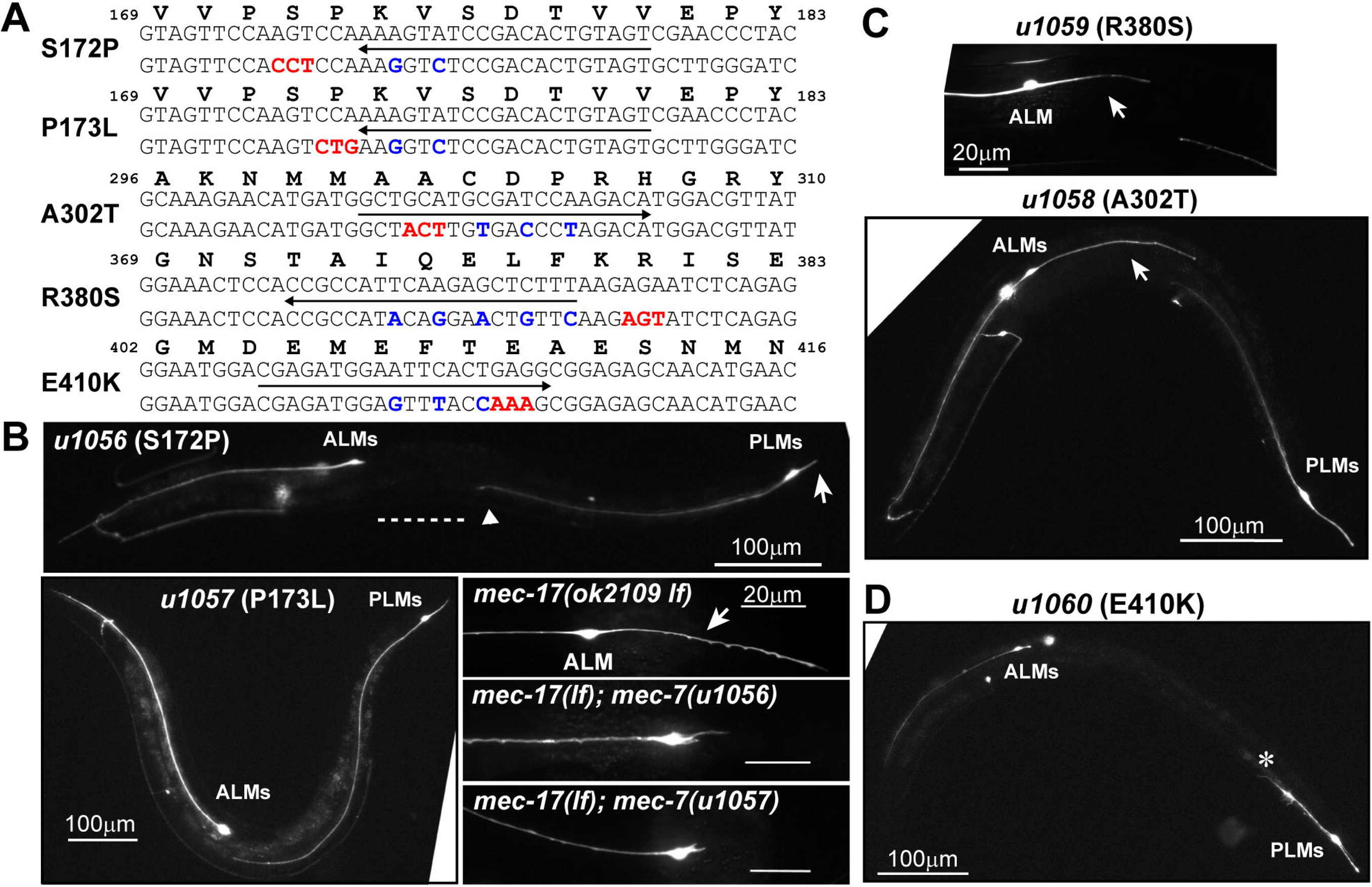
Modeling human tubulin mutations in *C. elegans* TRNs. (A) Sequence changes designed to introduce TUBB3 missense mutations in *mec-7* gene. For each mutation, the line with an arrow underscores the 20 bp DNA sequence for the guide RNA target, and the directionality of the arrow indicates the strand of the target. The bottom DNA sequence contains the desired nonsynonymous (red) and synonymous (blue) mutations on the homologous repair template. Amino acids sequence with the positional information was shown on the top. (B-D) TRN morphologies of animals carrying the engineered missense mutations. For *u1056*, the dashed line, triangle, and arrow indicate the gap from PLM-AN to ALM cell body, the position of the vulva, and the shortened PLM-PN, respectively. *u1057* showed normal TRN morphology. The growth of ALM-PN (arrow) was suppressed in *mec-17(ok2109 lf); mec-7(u1056)* and *mec-17(ok2109 lf); mec-7(u1057)* double mutants. For *u1058* and *u1059* (C), arrows point to the ectopic ALM-PN. For *u1060* (D), the asterisk demarcates the end of the severely shortened PLM-AN.

Heterozygous S172P and P173L mutations in human TUBB3 result in microcephaly, polymicrogyria, cortical dysplasia, and agenesis of the corpus callosum, all of which are defects in neuronal migration and axon growth (Jaglin *et al.*, 2009; Poirier *et al.*, 2010; Bahi-Buisson *et al.*, 2014). *In vitro* studies found that tubulin heterodimer containing the β-tubulin S172P mutant could not be incorporated into MTs (Jaglin *et al.*, 2009), indicating that the mutated protein is nonfunctional. Consistent with these findings, the *u1056* (S172P) mutation of *mec-7* led to a recessive phenotype similar to *mec-7(lf)* in the TRNs, which showed significantly shortened PLM-PN and slightly shortened PLM-AN with branching defects (Figure 5B). Surprisingly, the *u1057* (P173L) allele did not cause any TRN morphological or functional defects (Figure 5B). The difference between the mutant phenotypes in humans and worms is puzzling, since both S172 and P173 are located on the GTP binding B5-to-H5 loop (Figure 3C), which is identical in *C. elegans* MEC-7 and human TUBB3 (Figure S6). Perhaps the differences arise from differences in MT structure and organization: the *C. elegans* TRNs have 15-p MTs that are bundled, whereas human neurons have 13-p MTs that are not bundled. In fact, when 15-p MTs were converted to 13-p MTs and the bundle was disrupted in *mec-17* (α-tubulin acetyl-transferase) mutants (Topalidou *et al.*, 2012), both MEC-7 S172P and P173L mutations led to decreased MT stability and the loss of ALM-PN (Figure 5B and S7). This result suggests that P173L mutation indeed compromises MT stability in TRNs under sensitized conditions.

Heterozygous A302T and R380C mutations in human TUBB3, which affect residues on the external surface of the MTs, were associated with moderate congenital fibrosis of the extraocular muscles 3 (CEFOM3), anterior commissure hypoplasia, and corpus callosum hypoplasia. Mutation of A302 and R380 in yeast led to the formation of highly stable, benomyl-resistant MTs (Tischfield *et al.*, 2010). Consistent with these observations, we found that the same mutations of MEC-7 produced a *neo* phenotype (the generation of an ectopic ALM-PN; Figure 5C).

TUBB3 E410K mutation was also found in patients with severe CEFOM3 and hypoplasia of anterior commissure and corpus callosum, but these patients also suffer from facial weakness and progressive axonal sensorimotor polyneuropathy (Tischfield *et al.*, 2010). In yeast, MTs containing β-tubulin with the E410K substitution were less stable and less resistant to benomyl and had markedly decreased plus-end accumulation of kinesin-like motor proteins, compared to MTs with A302T and R380C mutations (Tischfield *et al.*, 2010). In *C. elegans*, MEC-7(E410K) produced a distinct neurite growth phenotype: ALM-AN and PLM-AN, but not PLM-PN were significantly shortened, which indicates reduced MT stability; but the PLM-PN was not affected, making the phenotype also different from that of the *mec-7(anti)* alleles, which cause general defects in MT polymerization (Figure 5D). Since E410 is also exposed to the exterior of MTs like A302 and R380 (Figure 3E), these results suggest that the cellular impact of the missense mutations would depend on the specific amino acid change instead of the location of the affected residues alone.

### The loss of *tba-7/*α-tubulin also caused excessive posterior neurite growth

In addition to MEC-7 and MEC-12, we identified a second α-tubulin gene *tba-7* that also regulates neurite growth in the TRNs. In the course of a mutagenesis for mutants with abnormal TRNs (Supplemental Results), we isolated a missense *tba-7* allele *u1015* (G92D), which caused the growth of an ectopic ALM posterior neurite, resembling the phenotype of *mec-7* and *mec-12 neo* alleles (Figure 6A and B). This *tba-7* allele is likely a *lf* allele, because it failed to complement with *gk787939* (Q230*), which is presumably a null and produced a similar phenotype as *u1015* (Figure S1C). We also examined mutants of other tubulin isotypes, and did not find morphological defects in the TRNs (we did not test for redundancy among the genes; Table S2).

**Figure 6.**
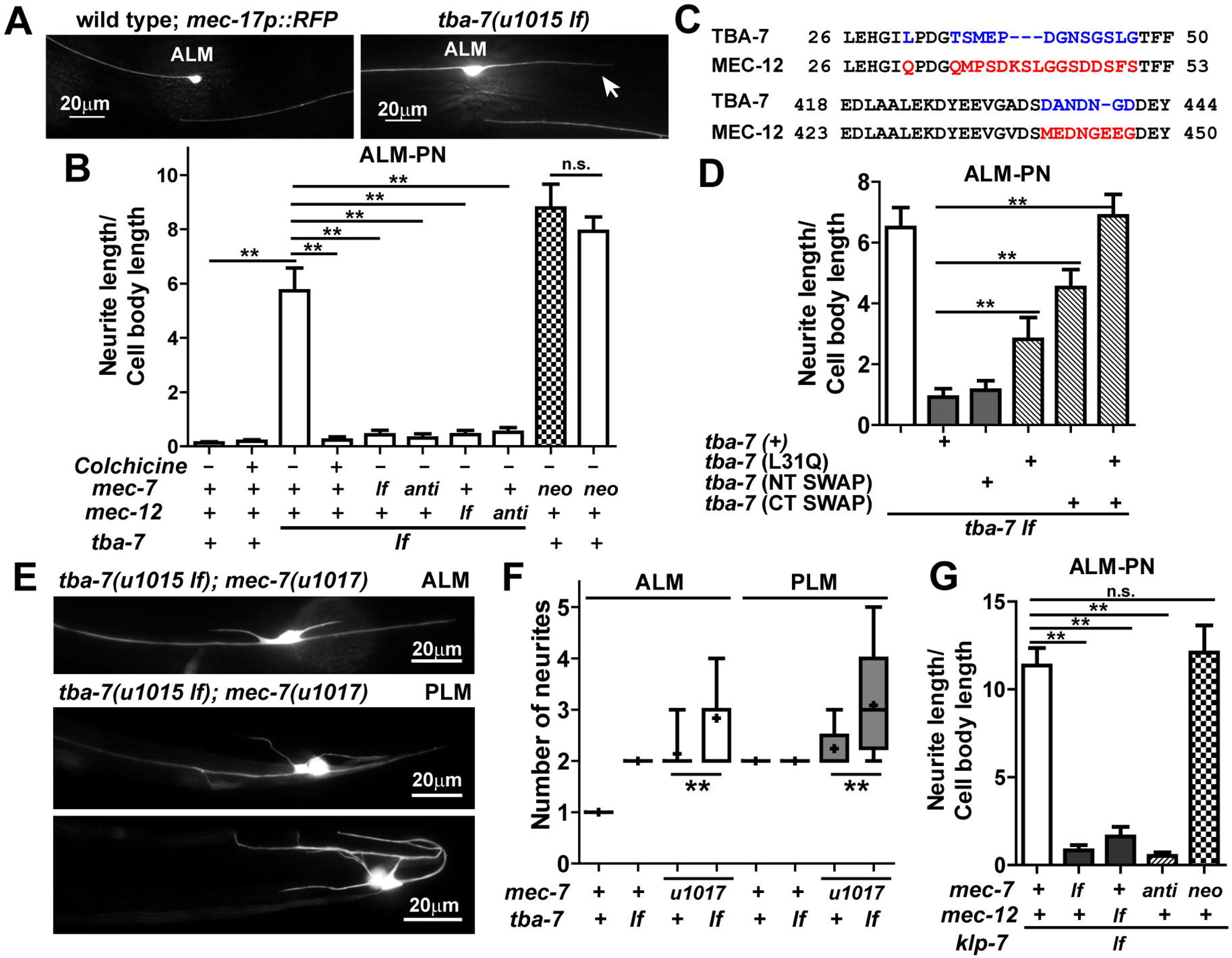
Mutations in *tba-7*, and *klp-7* result in the growth of ectopic ALM posterior neurites. (A) A representative image of the ectopic ALM-PN in *tba-7(u1015 lf)* mutants. (B) The length of ALM-PN in *tba-7(u1015 lf)* and the effect of 1 mM colchicine and *mec-7(ok2152 lf)*, *mec-7(u957 anti*), *mec-12(tm5083 lf)*, and *mec-12(u1016 anti)* mutations on the *tba-7(lf)* phenotype. (C) The differences between the TBA-7 and MEC-12 amino acid sequences in the alignment were labeled in color. (D) The length of ALM-PN in *tba-7(u1015 lf)* carrying the transgene expressing the wild type or mutant TBA-7 proteins. TBA-7 (NT SWAP) had TSMEPDGNSGSLG (a.a. 35-47) replaced by QMPSDKSLGGSDDSFS, TBA-7 (CT SWAP) had DANDNGD (a.a. 435-441) replaced by MEDNGEEG, and TBA-7 (L31Q + CT SWAP) carried both mutations. (E) Sprouting of neurites from the ALM and PLM cell bodies in *tba-7(u1015 lf)*; *mec-7(u1017 neo)* double mutants. (F) The number of neurites emanating from the cell bodies in *tba-7(u1015 lf)* and *mec-7(u1017 neo)* single mutants and their double mutants. Box plot shows the minimum, first quartile, median, third quartile, and maximum; the mean is indicated by a + sign. (G) The length of ALM-PN in *klp-7(tm2143 lf)* mutants and their double mutants with *mec-7(ok2152 lf)*, *mec-7(u957 anti*), *mec-12(tm5083 lf)*, and *mec-7(u1017 neo)*.

*tba-7* was expressed in the TRNs, and the excessive growth phenotype of *tba-7(u1015)* could be rescued by expressing *tba-7(+)* from the TRN-specific *mec-17* promoter, indicating that TBA-7 acts cell-autonomously in the TRNs (Figure S8). *tba-7*(*lf)* mutants did not exhibit touch insensitivity, synaptic vesicle mistargeting, or reduced protein levels, suggesting that the general function of TRN MTs was not affected and that the primary function of TBA-7 is to regulate neurite growth (Figure S9). We hypothesized that the loss of TBA-7 led to the formation of hyperstable MTs, similar to the *mec-7* and *mec-12 neo* mutations. Supporting this hypothesis, electron microscopy studies revealed that, like *mec-7(neo)* mutants, *tba-7(u1015)* animals retained the large diameter 15-p MTs and have closely packed MT bundles, which occupied a larger-than-normal area of a neurite cross-section (Figure 1). *tba-7(lf)* mutants also retained normal tubulin acetylation levels (Figure S9C) and had increased resistance to colchicine (Figure S3), which confirmed the presence of stable MTs. Treatment with 1 mM colchicine fully suppressed in the growth of ALM-PN in *tba-7(u1015)* animals, as did either *lf* or *anti* mutations in *mec-7* or *mec-12* (Figure 6B).

The amino acids sequences of α-tubulin TBA-7 and MEC-12 are 82% identical and 94% similar; they mainly differ in a N-terminal region and the C-terminal tail (Figure 6C). Domain swapping experiments showed that changing L31 of TBA-7 to Q found in MEC-12 or replacing the C-terminal DANDNGD (a.a. 435-441 in TBA-7) sequence with MEDNGEEG (a.a. 440-447 in MEC-12) partially impaired the function of TBA-7 proteins to restrict excessive neurite growth; and TBA-7 proteins with both L31Q mutation and the C-terminal replacement completely lost the ability to rescue the *tba-7(lf)* phenotype (Figure 6D). Q31 in MEC-12 is a potential site for polyamination (Song *et al.*, 2013) and E445 of the GEE motif (a.a. 444-446 in MEC-12, but absent in TBA-7) is a target for polyglutamination (Edde *et al.*, 1990). Post-translational modifications at these two sites could increase the stability of neuronal MTs (Song and Brady, 2015). Interestingly, replacing the a.a. 35-47 region of TBA-7 with the sequence from MEC-12 that contains lysine 40 (the site for α-tubulin acetylation) did not affect the TBA-7 function (Figure 6D), suggesting that the absence of this MT acetylation site in TBA-7 was not responsible for its activity in preventing ectopic neurite growth.

Since TBA-7 lacks some modification sites (e.g. Q31 and E445) that could stabilize MTs, our results suggest that TBA-7 may be a MT-destabilizing tubulin isotype, as compared to MEC-12. In wild-type animals, TBA-7 incorporation into the 15-p MTs with MEC-7 and MEC-12 may reduce MT stability; when TBA-7 is not available, more MEC-12 and possibly other α-tubulin isotypes replace TBA-7 and this replacement led to the formation of hyperstable MTs.

The *tba-7(u1015 lf); mec-7(u1017 neo)* double mutant had an ALM-PN that was similar in length to that in *mec-7(u1017)* single mutants (Figure 6A) but additionally produced up to 4 and 5 short ectopic neurites sprouting from the ALM and PLM cell bodies, respectively; this phenotype was rarely observed in *mec-7(u1017)* animals (Figure 6E and F). The additive effect in the double mutant suggest that the loss of TBA-7 and the MEC-7 *neo* mutations act in parallel to elevate MT stability.

### Mutations in MT-destabilizing Kinesin 13 affect TRN neurite growth

Since motor proteins and MAPs regulate MT dynamics (Akhmanova and Steinmetz, 2015), we expected mutations in MAP genes to cause similar phenotype to those of some of the tubulin mutations. Indeed, Ghosh-Roy *et al.* (2012) found and we have confirmed that the loss of the MT-depolymerizing kinesin *klp-7* induced the growth of an ectopic posterior neurite in ALM neurons (Figure 6G). KLP-7 belongs to the kinesin-13 family of catastrophe factors that bind to MT plus-ends and promotes depolymerization, thus generating dynamic microtubules (Han *et al.*, 2015). Importantly, we found that, similar to the tubulin *neo* alleles, the growth of the ectopic ALM-PN in *klp-7(lf)* mutants was suppressed by the loss of *mec-7* or *mec-12*, but the *klp-7* phenotype was not enhanced by *mec-7(neo)* mutations. These results suggest that the mutated MEC-7 proteins may render MTs hyperstable by reducing their interaction with KLP-7 or by making the MTs insensitive to the action of KLP-7.

## Discussion

### Neuronal morphogenesis requires optimal MT stability

MTs both support and regulate neurite growth. As the building block of MTs, α- and β-tubulins determine the structural properties of MTs and mediate their interaction with motor proteins and MAPs. As a result, any alteration in the tubulin proteins can potentially lead to changes in MT stability, which then affects neurite formation, guidance, and extension. In this study, we analyzed the effects of 67 tubulin missense mutations on neurite development in *C. elegans* touch receptor neurons. Based on the phenotypes, we categorized these mutations into three classes: loss-of-function (*lf*), antimorphic (*anti*), and neomorphic (*neo*) mutations. 1) *lf* mutations cause only mild neurite growth defects by reducing MT stability moderately; 2) *anti* mutations block MT polymerization and cause severe neurite growth defects; 3) *neo* mutations cause excessive neurite growth by inducing hyperstable MTs (Figure 7). The directionality of the excessive neurite in ALM reflects its potential to grow towards the posterior. In fact, about 30% of the adult ALM neurons have a very short (less than one cell body length) posterior protrusion, and manipulation of cellular signaling (e.g. activating the small GTPase Rac) can induce the production of ALM-PN (Zheng *et al.*, 2016). Thus, the ectopic growth of ALM-PN may serve as an indication for altered cytoskeletal organization and dynamics.

**Figure 7.**
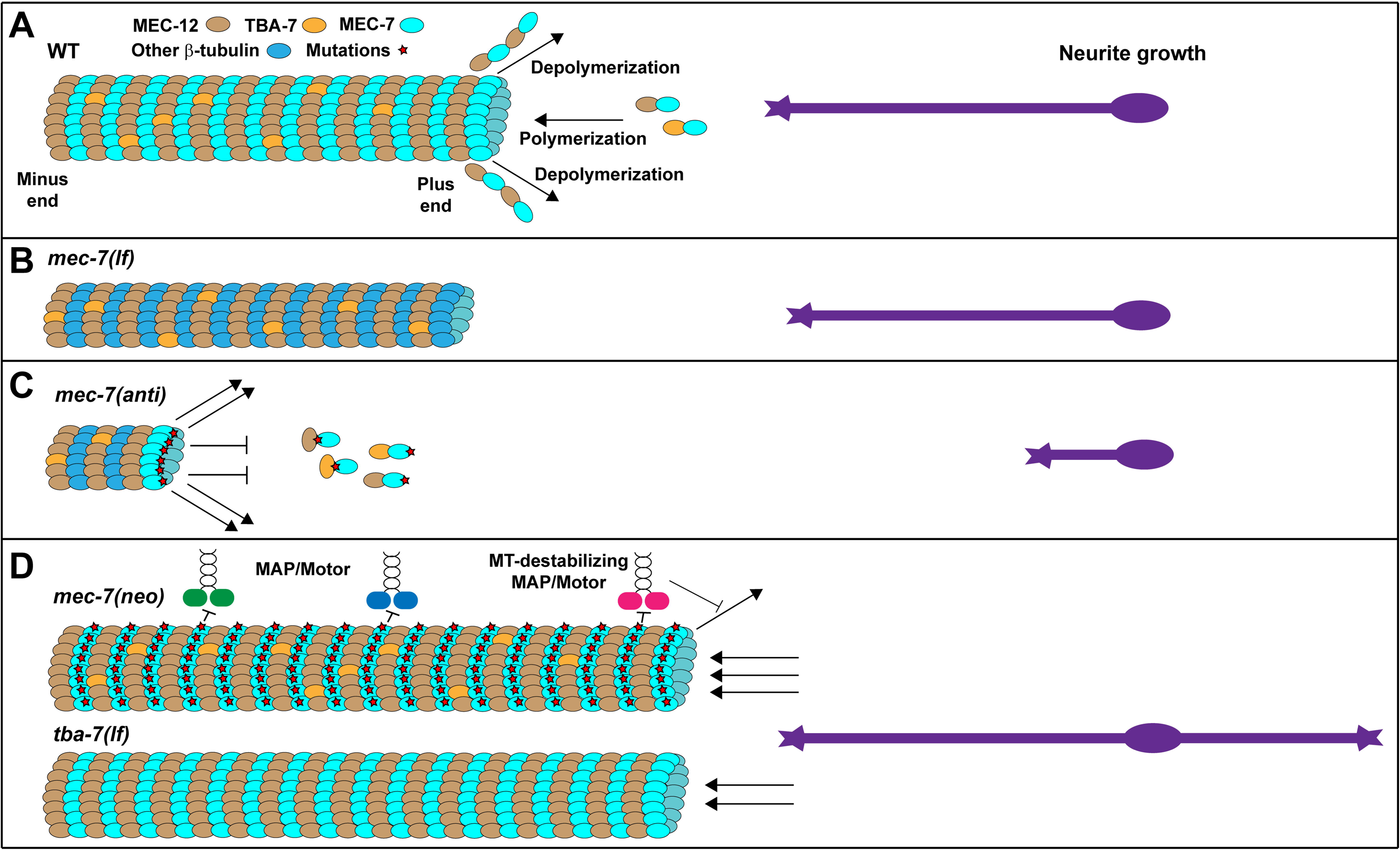
A model for the effect of MT dynamics on neurite development. (A) In wild-type animals, 15-p MTs are mostly made of MEC-12/α-tubulin (brown) and MEC-7/β-tubulin (cyan). Another α-tubulin TBA-7 (yellow) is also incorporated into the MTs, probably to decrease the stability. For simplicity, other tubulin isotypes that may function in TRNs, including TBA-1, TBA-2, TBB-1, and TBB-2 (Lockhead *et al.*, 2016), were shown in blue. Arrows towards the left and right represent depolymerization and polymerization, respectively. Stars represent mutations on the tubulin protein. (B) In the absence of MEC-7, the abundant 15-p MTs are replaced by a few 11-p MTs, which use other β-tubulins for polymerization and can still support neurite growth. Whether those β-tubulins are normally incorporated into the MTs is unclear. (C) *mec-7(anti)* mutations produce dominant-negative MEC-7 proteins with defects in the GTP binding pocket or the intradimer or interdimer interfaces. Such MEC-7 mutants can either sequester all α-tubulin or form toxic heterodimers that can terminate MT elongation upon incorporation. Thus, MT polymerization is blocked and neurite growth is disrupted. (D) *mec-7(neo)* mutations mostly map to the exterior facing surface of the tubulin structure. When incorporated into the MTs, those MEC-7 mutant proteins may affect the binding of MAPs or motor proteins (e.g. Kinesin 13 family protein KLP-7) by either binding poorer to destabilizing proteins or better to stabilizing proteins. The net result would be hyperstable MTs. The loss of TBA-7 presumably allowed more MEC-12 to be incorporated into the MTs, which led to increased MT stability. Abnormally elevated MT stability caused excessive neurite growth.

The correlation between the amount of neurite growth and the stability of MTs suggests that MT structure and dynamics are major determinants of neurite growth. Given the great number of proteins involved in both positively and negatively regulating MT assembly and disassembly (Mimori-Kiyosue, 2011), our results support the hypothesis that an optimal level of MT stability is required for proper neuronal morphogenesis. The finding that negative regulators of MT stability, such as the α-tubulin isotype TBA-7 and the MT-destabilizing kinesin-13 KLP-7, prevent excessive TRN neurite growth further supports this hypothesis.

### Structure-function analysis of tubulin mutations

The large number of missense mutations analyzed in this study allowed us to perform a structure-function analysis of neuronal tubulin proteins in living animals. Mapping the altered amino acid residues onto the *Bos taurus* tubulin α/β heterodimer structure 1JFF (Nogales *et al.*, 1998), we found that *lf* mutations mostly affect residues located in the interior of the structure and possibly disrupt protein folding. *anti* mutations affected residues that participate in GTP binding, intradimer interaction, or interdimer interactions. *neo* mutations mostly altered residues on the exterior surface and probably perturb the association of motor proteins and MAPs with MTs (Figure 3 and 7).

Our findings are consistent with similar structure-function relationship studies conducted in yeast α-tubulin *TUB1* and β-tubulin *TUB2* and Drosophila testis-specific *β2*-tubulin (Reijo *et al.*, 1994; Fackenthal *et al.*, 1995; Richards *et al.*, 2000) and provide another model to study the properties of tubulin proteins. For example, yeast *TUB1* mutations that caused supersensitivity to the MT-destabilizing drug benomyl mainly altered amino acids involved in GTP binding (e.g. D70) and intradimer (e.g. E98) and interdimer interaction; the change of equivalent residues in MEC-12 (D69 and E97) were identified in *mec-12(anti)* mutants. Similarly, some Drosophila *β2*-tubulin mutations (S25L and G96E) that made MTs less stable (Fackenthal *et al.*, 1995) affected the same or the adjacent residue in *mec-7(anti)* alleles [*u319* (S25F) and *u430* (A97V)].

Moreover, yeast *TUB1* mutations conferring benomyl-resistance mostly affected the residues on the outer surface of the MTs (Richards *et al.*, 2000) as did the *mec-7(neo)* mutations similarly that were resistant to colchicine. Thus, comparative studies using the three different organisms can help identify key residues in the tubulin structure responsible for various MT functions.

### Modeling the effects of neuronal tubulin mutations

We have used the *C. elegans* TRNs as a model to study neuronal MTs, which are much more stable and have different types of post-translational modification than the MTs in dividing cells (Baas *et al.*, 2016). Clinical studies in the past five years identified more than 100 missense mutations in tubulin genes that cause a wide spectrum of neurodevelopmental disorders, such as microcephaly, lissencephaly, polymicrogyria, and cortical malformation, along with severe defects in axon guidance and growth, including the complete or partial absence of corpus callosum, defects in commissural fiber tracts, and degeneration of motor and sensory axons (Tischfield *et al.*, 2011; Chakraborti *et al.*, 2016). Patients carrying different tubulin mutations often display distinct symptoms, but the complexity of the human nervous system makes it difficult to analyze the impact of specific mutations at the cellular level.

Combining CRISPR/Cas9-mediated genome editing with *in vivo* examination of TRN morphology, we were able to create several disease-causing human β-tubulin mutations in the *C. elegans mec-7* gene and examined their effects on neurite growth at a single neuron resolution. Our phenotypic evaluation was largely consistent with previous characterization of those mutations, suggesting that the TRN system could be instrumental to model the effects of human tubulin mutations identified in the clinic.

In fact, many disease-causing tubulin mutations alter amino acids that are physically adjacent to or located in the same region as the ones affected in our mutants, suggesting that they may affect MT stability in similar ways. By mapping 51 TUBA1A, 24 TUBB2B, and 19 TUBB3B mutations that were clinically identified onto the tubulin structure, we could correlate the location of some of the affected residues with the resulting clinical manifestation (Table S1). For example, 7/8 (87.5%) mutations that alter residues involved in GTP binding caused complete agenesis of corpus callosum, an indication of severe defects in axonal growth, whereas only 5/23 (21.7%) changes of residues in the interior of the structure and 10/32 (31.2%) mutations of residues involved in MAP binding did so (Table S1).

Position of the mutated residues alone, however, could not determine the effects of the missense mutation. For example, although the β-tubulin B5-to-H5 loop (from V170 to V179) contacts GTP, and four strong *mec-7(anti)* mutations (P171S, P171L, S176F, and V179A) mapped to this loop, the S172P mutation led to a *lf* phenotype and P173L mutation caused no defects in TRNs. One possible explanation is that S172 and P173 are further away from the GTP binding site, thus their mutations interfere less with microtubule function.

The identity of the residue replacing the wild type one is also important. For instance, *u278* (C303Y) was the strongest *mec-7(neo)* mutant, but *gk286001* (C303S) had no effect at all; since both tyrosine and serine have the polar hydroxyl group, the bulky phenyl group of the tyrosine may be responsible for disrupting some MT functions. Moreover, the *u1058* (A302T) mutation also caused a strong *mec-7(neo)* phenotype. A302 and C303 on the H9-to-B8 loop are exposed to the exterior of MTs but are not located on the major landing surface (H11 and H12) for motor protein and MAPs (Nogales *et al.*, 1998). Our results suggest that A302 and C303 may represent a previously unrecognized site of interaction between the MTs and their associated proteins. Even for residues located on H11 and H12, the effects of their mutation can be different. *u1059* (R380S) is a *mec-7(neo)* mutation, whereas *u1060* (E410K) allele produced an *anti*-like phenotype, suggesting that the impact of the missense mutation depends on how the interaction of MTs with the motor proteins or MAP is affected by the amino acid change.

### MT-destabilizing tubulin isotype and “multi-tubulin hypothesis”

The presence of multiple tubulin genes in eukaryotic genomes, their subtle differences in the protein sequences, and their functional differences led to the “multi-tubulin hypothesis.” In multicellular organism, the tubulins may have distinct expression patterns, *e.g.* Drosophila *β2*-tubulin is only expressed in the male germline (Kemphues *et al.*, 1980); human TUBB3B is highly enriched in the nervous system, whereas TUBB2B is expressed in many tissues (Sullivan and Cleveland, 1986). In single-cell organisms like *Tetrahymena*, different β-tubulin isotypes are spatially separated; some are only used to make the nuclear MTs of the mitotic apparatus, and some are only detected in the MTs of somatic cilia and basal bodies (Pucciarelli *et al.*, 2012). Our studies, however, found that two distinct α-tubulin isotypes (MEC-12 and TBA-7) could have different functions when both incorporated into the same MTs in the same neurons; MEC-12 promotes stability and TBA-7 promotes instability.

The abundance of large diameter 15-p MTs in the TRNs requires MEC-12, but the role of TBA-7 is to increase MT dynamics and to prevent the ectopic growth of posteriorly directed neurites in ALMs. Thus, TBA-7 is a MT-destabilizing tubulin isotype, and this destabilizing activity depends on the absence of potential polyamination and polyglutamination sites that could increase MT stability. Our results are consistent with early *in vitro* experiments suggesting that TUBB3 (also known as βIII) also promotes MT dynamics; removal of *T*U*BB3* from the brain extract resulted in a tubulin mixture that assembled much more rapidly than the unfractionated control (Banerjee *et al.*, 1990), and MTs assembled from the purified *αβ*_III_ heterodimers were considerably more dynamic than MTs made from the *αβ*_II_ and *αβ*_IV_ dimers (Panda *et al.*, 1994).

However, the long-standing question is whether some tubulin isotypes do destabilize MTs *in vivo*. Our findings suggest the answer is yes. A balanced incorporation of multiple tubulin isotypes into the same MT structure is critical to generate MTs with the optimal stability.

The absence of post-translational modification sites in tubulin isotypes may lead to functional diversity. In addition to TBA-7, among the other eight *C. elegans α*-tubulin isotypes, TBA-8 and TBA-5 also lack the Q31 for polyamination, and TBA-6 lacks the E445 for polyglutamination; all nine human α-tubulins have Q31, but TUBAL3 has a short C-terminal tail and does not contain potential polyglutamination sites (Figure S10). The incorporation of those tubulin isotypes may be a general regulatory mechanism to control MT stability. Despite the functional differences between MEC-12 and TBA-7, the fact that both *mec-12(lf)* mutants and *mec-12(lf); tba-7(lf)* double mutants retained the ability to grow out a normal amount of neurites suggests also genetic redundancy among the α-tubulins in supporting neurite growth.

Different contributions of tubulin isotypes to MT dynamics were also observed in *C. elegans* embryogenesis (Honda *et al.*, 2017). The spindle MTs required two β-tubulins (TBB-1 and TBB-2), but TBB-2 was incorporated into the MTs twice as much as TBB-1; the loss of TBB-2 caused a dramatic decrease in MT growth rate and an increase in catastrophe frequency, leading to highly unstable MTs, whereas the loss of TBB-1 only slightly reduced growth rate. These data support the “multi-tubulin concept” that different tubulin isotypes have distinct functions in the same cells.

## Materials and Methods

### Strains and genetic screens

*C. elegans* wild type (N2) and mutant strains were maintained at 20 °C as previously described (Brenner, 1974). *mec-7* alleles *u910*, *u911, u955*, *u956*, *u957*, *u958*, *u1017*, and *u1020*, *mec-12* alleles *u917*, *u950*, *u1016*, *u1019*, and *u1021*, and *tba-7(u1015)* were isolated by visually screening for mutants with TRN differentiation defects using TU4069, which carries *uIs134 [mec-17p::RFP]* for the visualization of the TRNs, as the starter strain and ethyl methanesulfonate as the mutagen (Brenner, 1974). Mutants were outcrossed with wild type, and the phenotype-causing mutations were identified as previously described by whole-genome resequencing (Zheng *et al.*, 2013) or by complementation tests with reference alleles. A detailed description of the screen can be found in the supplemental results.

*mec-7* alleles *e1343*, *e1505*, *e1527*, *e1522*, *n434*, *u10*, *u18*, *u22*, *u48*, *u58*, *u98*, *u127*, *u129*, *u162*, *u170*, *u223*, *u225*, *u278*, *u234*, *u249*, *u262*, *u275*, *u283*, *u305*, *u319*, *u428*, *u429*, *u430*, *u433*, *u449*, and *u445*, and *mec-12* alleles *e1605*, *e1607*, *u50*, *u63*, *u76*, and *u241* were previously isolated (Chalfie and Sulston, 1981; Chalfie and Au, 1989; Savage *et al.*, 1994) and re-examined in this study.

*u1056* to *u1060* alleles were created through CRISPR/Cas9-mediated homologous recombination (Dickinson *et al.*, 2013). Guide RNAs were designed according to the target sequence closest to the desired codon change (Figure 5A). Recombination templates were created by cloning the *mec-7* coding region and 466 bp 3’UTR sequence into the SalI and BamHI sites of pBlueScript II SK (+) vector and then introducing the desired missense mutation (red in Figure 5A), along with synonymous mutations (blue in Figure 5A) that change the guide RNA target sites, using the Q5 Site-Directed Mutagenesis Kit from New England Biolabs (NEB; Ipswich, MA). pDD162 constructs expressing Cas9 and specific guide RNAs were injected together with the corresponding recombination template and marker *myo-2p::RFP* into TU4069 animals. F1s expressing RFP in the muscle were singled out and genotyped to identify heterozygotes with successful edits, and F2s carrying homozygous mutations were then isolated and examined for TRN morphology.

The *gk* alleles of *mec-7* and *mec-12* listed in Table 1 and *tba-7(gk787939)* were generated in the million mutation project (Thompson *et al.*, 2013). These *gk* alleles, as well as *mec-17(ok2019)*, *dlk-1(ju476)*, and *dlk-1(km12)* were obtained from the Caenorhabditis Genetics Center, which is funded by NIH Office of Research Infrastructure Programs (P40 OD010440). *klp-7(tm2143)*, and *mec-12(tm5083)* were generated by the National Bioresource Project of Japan, and *mec-12(gm379)* was kindly provided by Dr. Chun-Liang Pan at the National Taiwan University.

Additional *mec-12* null alleles (*u1026*, *u1027*, and *u1028*) were generated by CRISPR/Cas9-mediated genome editing targeting 5’-GAAGTAATTTCGATTCACATCGG-3’ in exon 2 of *mec-12* as described above (Dickinson *et al.*, 2013). Frameshift-causing mutations were identified by sequencing the *mec-12* locus (Figure S2).

### Constructs and Transgenes

A *tba-7::GFP* reporter (TU#1632) was made by cloning a 1.2 kb *tba-7* promoter and the entire coding region from the genomic DNA to wild-type genomic DNA into the Gateway pDONR221 P4-P1r vector; the resulting entry vector, together with the pENTR-GFP, pENTR-*unc-54*-3’UTR, and destination vector pDEST-R4-R3 were used in the LR reaction to create the final expression vectors. *mec-17p::tba-7(+)* (TU#1629) was created by cloning a 1.9 kb *mec-17* promoter and the *tba-7* coding sequence into pDONR221 P4-P1r and pDONR221 vectors, respectively, and assembling these entry vectors with pENTR-*unc-54*-3’UTR and the destination vector. Details about the Gateway cloning method by Life Technologies can be found at www.invitrogen.com/site/us/en/home/Products-andServices/Applications/Cloning/Gateway-Cloning.html. *tba-7(+)* locus, including the 1.2 kb promoter, coding region, and a 911-bp 3’UTR, was cloned using SalI and NotI into pBlueScript II SK(+) vector; NEB Q5 site-directed mutagenesis kit was then used to generate constructs expressing the TBA-7 variants shown in Figure 6E.

Transgenes *uIs31[mec-17p::GFP]* III, *uIs115[mec-17p::RFP]* IV, and *uIs134[mec-17p::RFP]* V were used to visualize TRN morphology (Zheng *et al.*, 2015). *jsIs821[mec-7p::GFP::RAB-3]* was used to assess synaptic vesicle localization (Bounoutas *et al.*, 2009b).

### Electron microscopy

We re-analyzed previously collected cross-section images of ALM neurites from wild type animals (N501, N631, N933, and N934 from the Hall lab collection) and from *mec-7(neo)* mutants *u170* and *u278* (Savage *et al.*, 1994). The wild type animals had been fixed by either chemical immersion fixation without tannic acid (samples N501, N631; Hall, 1995) or by high-pressure freezing/freeze substitution (HPF/FS) including tannic acid (samples N933, N934; Topalidou *et al.*, 2012). The *mec-7(neo)* alleles had also been fixed by chemical immersion, without tannic acid.

For this study, we fixed *mec-7(ok2152 lf)*, *mec-12(tm5083 lf)*, and *tba-7(u1015 lf)* adults using an HPF/FS protocol that included a first fixation in 0.5% glutaraldehyde + 0.1% tannic acid and a second fix in 2% osmium tetroxide + 0.1% uranyl acetate (UAc), both at -90 °C, followed by staining in 1% UAc in acetone at 0 °C. We also fixed adults of *mec-12(anti)* mutants, *u950*, *u76*, and *u1021*, using HPF/FS using a similar protocol, but without tannic acid. Eighty-nanometer transverse sections were collected at multiple positions along the ALM anterior neurite and post-stained with uranium acetate and lead citrate. A Philips CM10 electron microscope with an attached Morada digital camera (Olympus) was used to acquire the images.

For each animal, we counted the number of MTs in the ALMs and measured the MT diameter on sections collected from at least five different positions along the ALM-AN; data from ALML and ALMR were combined. For each strain, we examined sections from at three different animals. For wild-type animals, *tba-7(lf)*, and *mec-7(neo)* mutants, we used at least ten sections to measure the distance between two closest MT centers and the proportion of MT-occupied area to the cross-sectional area of the neurite.

We noticed that our earlier studies found that wild-type TRN MTs had much larger diameters (29.6 ± 0.4 nm; Chalfie and Thomson, 1979) than we find here (21.5 ± 1.6 nm). Fixation method could not explain the difference, because we obtained similar results from N501 and N631 (fixed by chemical immersion) and N933 and N934 (fixed by HPF/FS); data obtained using both methods showed smaller MT diameters than the measurements from earlier studies, which used chemical immersion fixation. Although we could not explain this discrepancy, we are very confident in the current calibration of the electron microscope.

### Phenotype Scoring and Statistical Analysis

We measured the length of TRN neurites in at least 30 fourth stage larvae or young adults grown at 20° C, except where otherwise stated. Relative length of posteriorly directed neurites was calculated by dividing the neurite length by the diameter of the cell body. Defects in TRN anterior neurite length were assessed by counting the percentage of cells whose neurites failed to reach the vulva (for PLM) or the posterior pharyngeal bulb (for ALM); at least 50 cells were examined for each strain.

Fluorescence intensity of the RFP expressed from the *uIs134* transgene in the ALM cell body was used to measure TRN protein levels; intensity was calculated after manual background subtraction using ImageJ as previously described (Chen and Chalfie, 2015). Synaptic vesicle localization was quantified by calculating the percentage of animals in each of the four phenotypical categories: 1) normal localization to the PLM synaptic branch; 2) reduced intensity of the synaptic marker at the synapses as partial defect; 3) complete loss of normal localization to the synapse as complete defect; 4) localization to the PLM-PN as mistargeting. At least 50 adult animals were examined.

Acetylated α-tubulin staining using antibody [6-11B-1] (Abcam, Cambridge, MA) was performed and analyzed as previously described (Topalidou *et al.*, 2012). For colchicine treatment, animals were grown in standard NGM agar plates containing different concentrations (from 0.06 mM to 2 mM) of colchicine before phenotypic analysis as described before (Bounoutas *et al.*, 2009a). Similarly, 100 nM paclitaxel was added to the NGM agar plates, L1 animals were placed onto the plates, and young adults were examined for TRN morphology. PyMOL (Schrodinger, 2015) was used to view the structure of α/β tubulin dimer (1jff.pdb; Nogales *et al.*, 1998) and to label affected residues in *mec-7* and *mec-12* mutants,.

For statistical analysis, ANOVA and the post hoc Dunnett’s test (comparison of each of many treatments with a single control) or Tukey-Kramer test (studentized range test for all pairwise comparison) were performed using the GraphPad Prism (version 5.00 for Windows, GraphPad Software, La Jolla California USA, www.graphpad.com) to identify significant difference between the mutants and wild type animals. Student’s *t* test was used to find significant difference between two samples in paired comparisons. Single and double asterisks indicated *p* < 0.05 and *p* < 0.01, respectively. The χ^2^ test was used for the categorical data to find significant difference between different strains.

## Acknowledgements

We thank Dan Dickinson, Chun-Liang Pan, and Andrew Chisholm for sharing reagents. This work was supported by NIH Grant GM30997 (to M.C.) and OD 010943 (to D.H.H.). Core

